# Biophysical investigation of the dual binding surfaces of human transcription factors FOXO4 and p53

**DOI:** 10.1101/2021.01.11.425814

**Authors:** Jinwoo Kim, Dabin Ahn, Chin-Ju Park

**Author notes:** These authors contributed equally to this work (Jinwoo Kim, Dabin Ahn).

## Abstract

Cellular senescence is protective against external oncogenic stress, but its accumulation causes aging-related diseases. Forkhead box O4 (FOXO4) and p53 are human transcription factors known to promote senescence by interacting each other and activating p21 transcription. Inhibition of the interaction is a strategy for inducing apoptosis of senescent cells, but the binding surfaces that mediate the FOXO4-p53 interaction remain elusive. Here, we investigated two binding sites involved in the interaction between FOXO4 and p53 by NMR spectroscopy. NMR chemical shift perturbation analysis showed that the binding between FOXO4’s forkhead domain (FHD) and p53’s transactivation domain (TAD), and between FOXO4’s C-terminal transactivation domain (CR3) and p53’s DNA binding domain (DBD), mediate the FOXO4-p53 interaction. Isothermal titration calorimetry data showed that both interactions have micromolar K_d_ values, and FOXO4 FHD-p53 TAD interaction has a higher binding affinity. Also, we showed that the FOXO4 CR3-binding surface of FOXO4 FHD interacts with p53 TAD2, and FOXO4 CR3 interacts with the DNA/p53 TAD-binding surface of p53 DBD, suggesting a network of potentially competitive and/or coordinated interactions. Based on the results, we propose that the dual interaction contributes to two TF’s proper location on the p21 promoter site and consequently promotes p21 transcription and cell senescence. This work provides structural information at the molecular level that is key to understanding the interplay of two proteins responsible for cellular senescence.

**Conflicts of interest:** *None*

## Introduction

The forkhead box class O (FOXO) subfamily consists of four transcription factors (TFs), FOXO1, FOXO3a, FOXO4, and FOXO6, which share a conserved DNA binding domain called the forkhead domain (FHD).[1, 2] As TFs, FOXO proteins participate in various cellular processes such as regulation of the cell cycle, cell survival, metabolism, and tumorigenesis by controlling the expression of their target genes.[3–10] In particular, FOXOs are known to be important regulators of cellular senescence.[11] Cellular senescence is defined as the induction of permanent cell cycle arrest regardless of the presence or activity of growth factors. Therefore, senescent cells avoid apoptosis and induce a pro-inflammatory environment that affects the neighboring cells.[12–14] Unlike apoptotic cells, which are immediately removed, senescent cells are maintained for a long time, and their accumulation accelerates aging and causes senescence-associated diseases such as type Ⅱ diabetes[15], osteoarthritis[16], atherosclerosis[17], neurodegenerative disorders[18], and cancers[13, 14, 19].

Like other FOXOs, FOXO4 has five domains: the FHD, a nuclear localization sequence, a nuclear export sequence, and two transactivation domains called CR1 and CR3 (Figure 1A).[20] The CR3 domain is also known as the transactivation domain (TAD). Only the FHD, which binds to two conserved DNA sequences, 5’-TTGTTTAC-3’, named the Daf-16 family member-binding element [21, 22], and 5’-(C/A)(A/C)AAA(C/T)AA -3’, named the insulin-responsive sequence [22], has a defined three-dimensional structure. A previous study suggested that FHD and CR3 interact to form an intramolecular complex, and CR3 kinetically affects selective DNA recognition of FHD.[23] In regions other than FHD, including CR3, the protein is intrinsically disordered.[24] After acute DNA damage, the expression level of FOXO4 gradually increases, which induces cellular senescence and suppresses the apoptosis response, allowing the viability of senescent cells to be maintained.[11] Inhibition of FOXO4 expression causes the release of mitochondrial cytochrome C and BAX/BAK-dependent caspase-3 cleavage, leading to reduced cell viability and density. Also, an increase in reactive oxygen species via oxidative stress triggers phosphorylation of c-Jun N-terminal kinase (JNK) through the mitogen‐activated protein kinase kinase pathway.[7] In turn, phosphorylated JNK phosphorylates and activates FOXO4.[6–8] Activated FOXO4 increases the transcription of p21, which is involved in cell cycle arrest and promotes cellular senescence.[7, 25]

**Figure 1.**
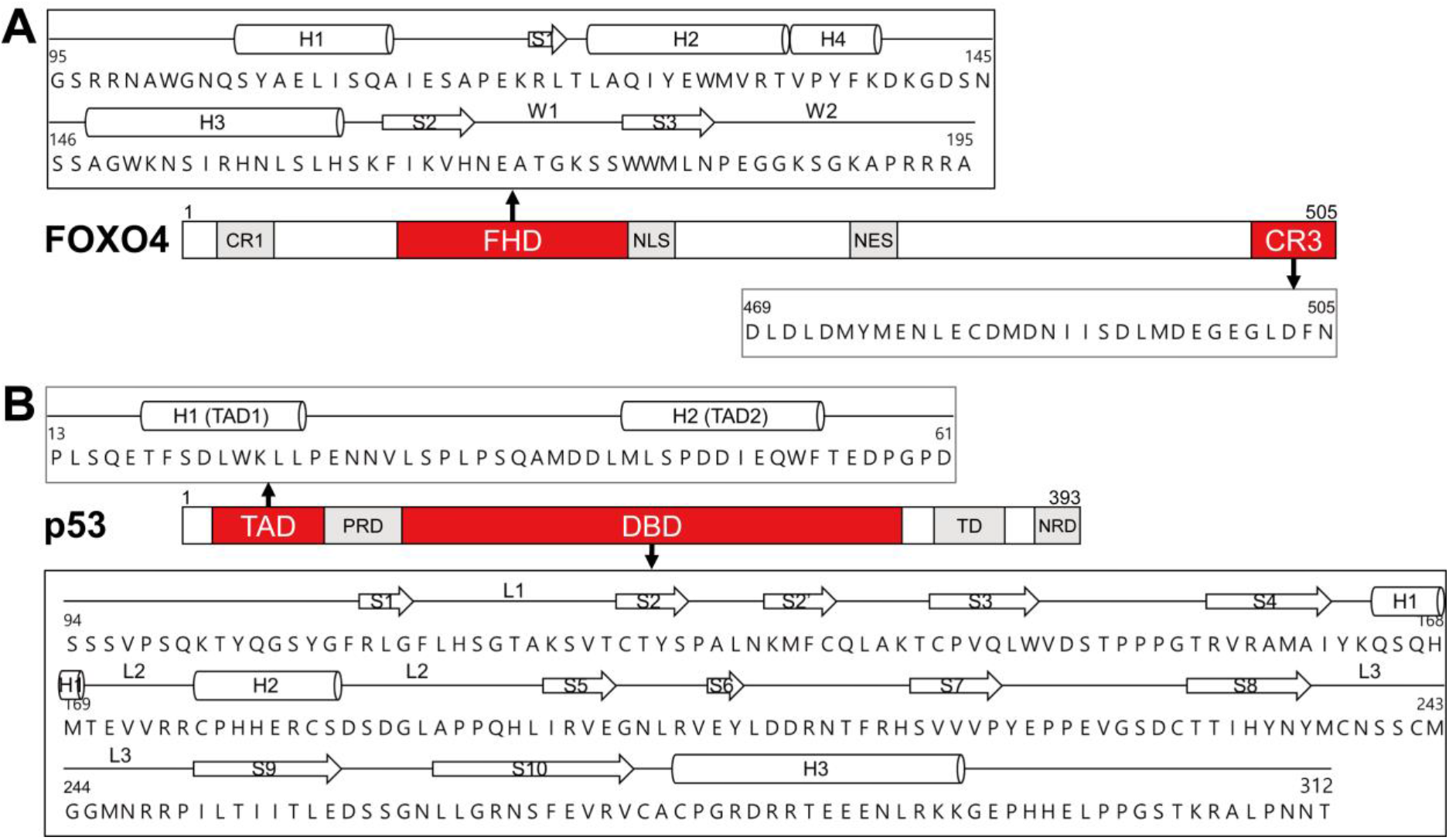
Domain structure of (A) FOXO4 and (B) p53. The primary sequences of FOXO4 FHD and CR3 and p53 TAD and DBD are shown as expansions indicated by arrows. CR1 and CR3, transactivation domains; DBD, DNA-binding domain; FHD, forkhead domain; NES, nuclear export sequence; NLS, nuclear localization sequence; NRD, negative regulatory domain; PRD, a proline-rich domain; TAD, transactivation domain; TD, tetramerization domain

As senescence progresses, FOXO4 is gradually accumulated into euchromatin foci and interacts with activated p53 in promyelocytic leukemia bodies at the DNA damage site.[11] P53 is a homo-tetrameric transcription factor that regulates various target genes and participates in the control of several cellular processes, including cell cycle arrest, apoptosis, and senescence.[26–30] The N-terminal transactivation domain (TAD), a proline-rich domain, a DNA-binding domain (DBD), a tetramerization domain (TA), and the C-terminal negative regulatory domain are components of p53 (Figure 1B). With these multiple domains, p53 can cooperate with numerous partners and behaves as a central protein-protein interaction (PPI) hub in biological systems.[31] Although p53 and FOXOs specifically recognize different sequences in promoter sites, they have common downstream target genes such as WIF1, FASLG, GADD45, and PA21.[32, 33] In particular, when FOXO4 and p53 bind each other within the nucleus of senescent cells, the nuclear exclusion of p53 is disrupted, and the transcription of the p21 gene is activated.[11, 34] Overexpressed p21 participates in the cellular senescence process by arresting the cell cycle through negative regulation of the G1/S and G2/M transitions.[35, 36] In addition, since the exclusion of p53 is restrained, the apoptosis process induced by p53 cannot proceed appropriately in mitochondria, accelerating cellular senescence.[37] Therefore, it has been proposed to modulate the FOXO4-p53 interaction as a target for treating aging. Baar *et al.* developed the FOXO4-D-retro-inverso (FOXO4-DRI) peptide as a way of introducing D-amino acids into a retro-reversed sequence of the FOXO4 FHD N-terminus.[11] They attempted to inhibit the binding between FOXO4 and p53 through FOXO4-DRI, and demonstrated that the introduction of this peptide induces death of senescent cells.

Despite the increasing interest in the FOXO4-p53 interaction, structural information of the complex is insufficient. Since it is known that the FOXO4 FHD binds a wide region of the p53 sequence (amino acids 1-312), including the TAD, the proline-rich domain, and the DBD,[11] it is necessary to clarify the precise binding site further to understand the FOXO4-p53 complex. Therefore, we investigated the interaction of FOXO4 and p53 at the molecular level. The FHD and CR3 domains of FOXO3a, which has high sequence similarity to FOXO4, bind to the DBD of p53.[38] Also, CR3 of the FOXO proteins plays a role in recruitment of co-activators such as CBP/p300.[38, 39] Therefore, we selected FHD and CR3 of FOXO4 as potential candidates for p53 binding. The DBD and TAD regions of p53 interact with many proteins and control various signaling pathways[38, 40–47], thus we examined these two domains as candidates for binding to FOXO4. In this work, we demonstrated that FOXO4’s FHD and CR3 directly bind to TAD and DBD of p53, respectively, using nuclear magnetic resonance (NMR) spectroscopy. Alanine mutation analysis of FOXO4 CR3 showed that the binding to p53 DBD occurs on a wide and shallow surface. A binding affinity comparison using isothermal titration calorimetry (ITC) suggested that FOXO4 CR3-p53 DBD binding takes precedence over FOXO4 FHD-p53 TAD during the formation of the FOXO4-p53 complex. Based on these findings, we proposed that the dual binding of FOXO4 and p53 could influence the formation of the cooperative p21 transcription initiation complex, which is necessary for maintaining cellular senescence. This biophysical characterization of the FOXO4-p53 interaction is expected to contribute to new strategies for developing inhibitors of cellular senescence.

## Results

### FOXO4 FHD prefers binding to p53 TAD

The binding of FOXO4 FHD to p53 was previously briefly studied using NMR spectroscopy.[11] However, it is still unclear which domain of p53 interacts with FOXO4 FHD and their mutual interfaces. In order to accurately identify the domain used for the interaction of p53 with FOXO4 FHD, we performed chemical shift perturbation (CSP) analysis of ^15^N-labeled FOXO4 FHD with unlabeled p53 TAD or DBD. CSP showed which domain of p53 was involved in the interaction with FOXO4 FHD when equimolar amounts of the respective domains were present. The ^1^H-^15^N amide cross-peaks of ^15^N-labeled FOXO4 FHD were assigned based on the previously published backbone assignment.[48] In the NMR titration between ^15^N-labeled FOXO4 FHD and p53 TAD, several residues were found to undergo a large change (Figure 2A and 2C). Residues G102, S105, and Y137 of FOXO4 FHD were perturbed by more than the average + 2 standard deviations (σ), and residues W101ε, Q104, Y106, L160, K163, T172, and E183 were also changed relatively significantly (average + 1σ). They are all located in a pocket-like space formed in the crevice between the N-terminal loop and α-helices H1, H4, and H3, except for T172, which is located in the flexible wing W1 (Figure 2E). These results are consistent with the previous titration data for p53 (1-312) and FOXO4 FHD.[11] They imply that p53 binds to FOXO4 FHD mainly through its TAD. Furthermore, W101ε, Q104, S105, L160, and T172 of FOXO4 FHD are residues that also participate in the intramolecular interaction of FOXO4 FHD with CR3.[49] This suggests that the same region of FOXO4 FHD is used for both intramolecular FOXO4 CR3 and intermolecular p53 TAD interactions. In addition, the large perturbation of three aromatic residues (Y137, W101ε, Y106) may mean that hydrophobic interactions are involved in the binding of FOXO4 FHD and p53 TAD.

**Figure 2.**
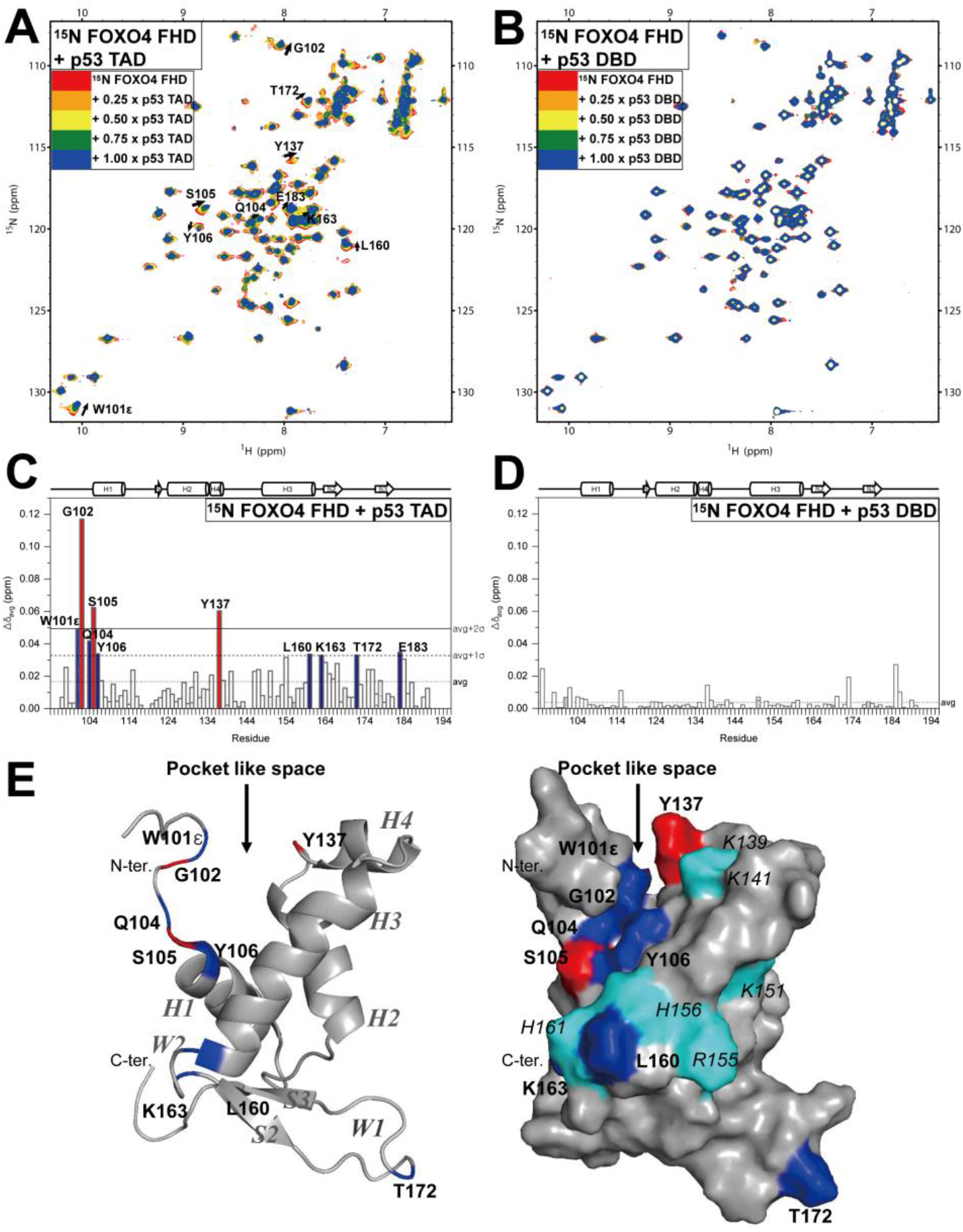
Chemical shift changes of FOXO4 FHD upon titration with p53 domains. (A-B) Overlaid ^1^H-^15^N HSQC spectra of ^15^N-labeled FOXO4 FHD at a concentration of 300 μM with increasing concentrations (0 μM, 75 μM, 150 μM, 225 μM, 300 μM) of (A) p53 TAD and (B) p53 DBD. (C-D) Δδ_avg_ for each FOXO4 FHD residue when (C) p53 TAD and (D) p53 DBD concentrations are 300 μM, respectively. (E) The solution structure of FOXO4 FHD (PDB ID: 1E17),[48] colored based on CSP upon addition of p53 TAD. Residues that undergo a large chemical shift change are colored in red (above average + 2σ) and blue (above average + 1σ). Basic amino acids of FOXO4 FHD are colored in cyan.

Next, we titrated unlabeled p53 DBD into ^15^N-labeled FOXO4 FHD (Figure 2B and 2D). In contrast to experiments with p53 TAD, we did not observe any significantly perturbed peaks of FOXO4 FHD upon p53 DBD titration. The average value of CSP of FOXO4 FHD caused by p53 DBD was about 0.004 ppm, which was much smaller than that caused by p53 TAD, 0.154 ppm. These results suggest that p53 interacts with FOXO4 FHD via its TAD domain rather than the DBD. It is expected that p53 TAD binds in the pocket provided by FOXO4 FHD, possibly interfering with the formation of the intramolecular complex of FOXO4 and inducing the FOXO4-p53 interaction.

### p53 helix TAD2 is responsible for the interaction with FOXO4 FHD

As shown in Figure 1B, the p53 TAD contains two α-helices, which are named TAD1 and TAD2. They are adjacent in the primary structure but seem to have distinct functions. For example, TAD1 is responsible for p53-dependent transactivation. It is essential for DNA damage-induced G1 arrest but not for RAS-induced senescence in fibroblasts,[50] while TAD2 predominates over TAD1 in binding to mediator complex subunit 25 (MED25).[51] However, in a p53 intramolecular interaction, both TAD1 and TAD2 participate in binding to p53 DBD.[52] In order to understand the interaction between FOXO4 FHD and p53 TAD more precisely, we studied which parts of p53 TAD are responsible for FOXO4 FHD binding. We titrated unlabeled FOXO4 FHD into ^15^N-labeled p53 TAD and performed the CSP analysis to observe whether TAD1 and TAD2 are cooperative or independent in FOXO4 binding.

In Figures 3A and 3C, the CSP of TAD1 and its nearby residues was relatively small. There were no residues that changed more than the average + 1σ. In contrast, TAD2 had a completely different trend. Residues D48, E51, Q52, W53, and F54 in TAD2 had perturbation of more than the average + 2σ, and it was the same for neighboring T55. I50 also underwent a relatively large change. These results strongly indicate that p53 TAD2 is dominantly involved in FOXO4 FHD binding, whereas TAD1 does not participate or has a minor role. Additionally, changes of A39, M40, D42, L43, L45, and S46 near TAD2’s N-terminus were observed, which may imply that not only TAD2 but also its surrounding residues participate in the interaction with FOXO4 FHD, or that their structural environment is changed by the interaction. As found in FOXO4 FHD, p53 TAD2 and its surroundings showed changes in aromatic residues (W53 and F54) and perturbation of several aliphatic residues (L32, A39, M40, M44, L45, I50) was observed. This underscores the hydrophobic portion of the interaction. It is also noteworthy that several acidic residues were perturbed in p53 TAD2 (D48, E51, and D57), which are complementary to the basic amino acids K139, K141, K151, R155, H156, and H161 located on the FOXO4 FHD H1, H3, and H4 binding surface (Figure 2E). Also, FOXO4 FHD is positively charged (pI = 10.11) and p53 TAD (pI = 3.39) is negatively charged. Thus, electrostatic contacts can be reasonably expected to contribute to the interaction between FOXO4 FHD and p53 TAD2.

**Figure 3.**
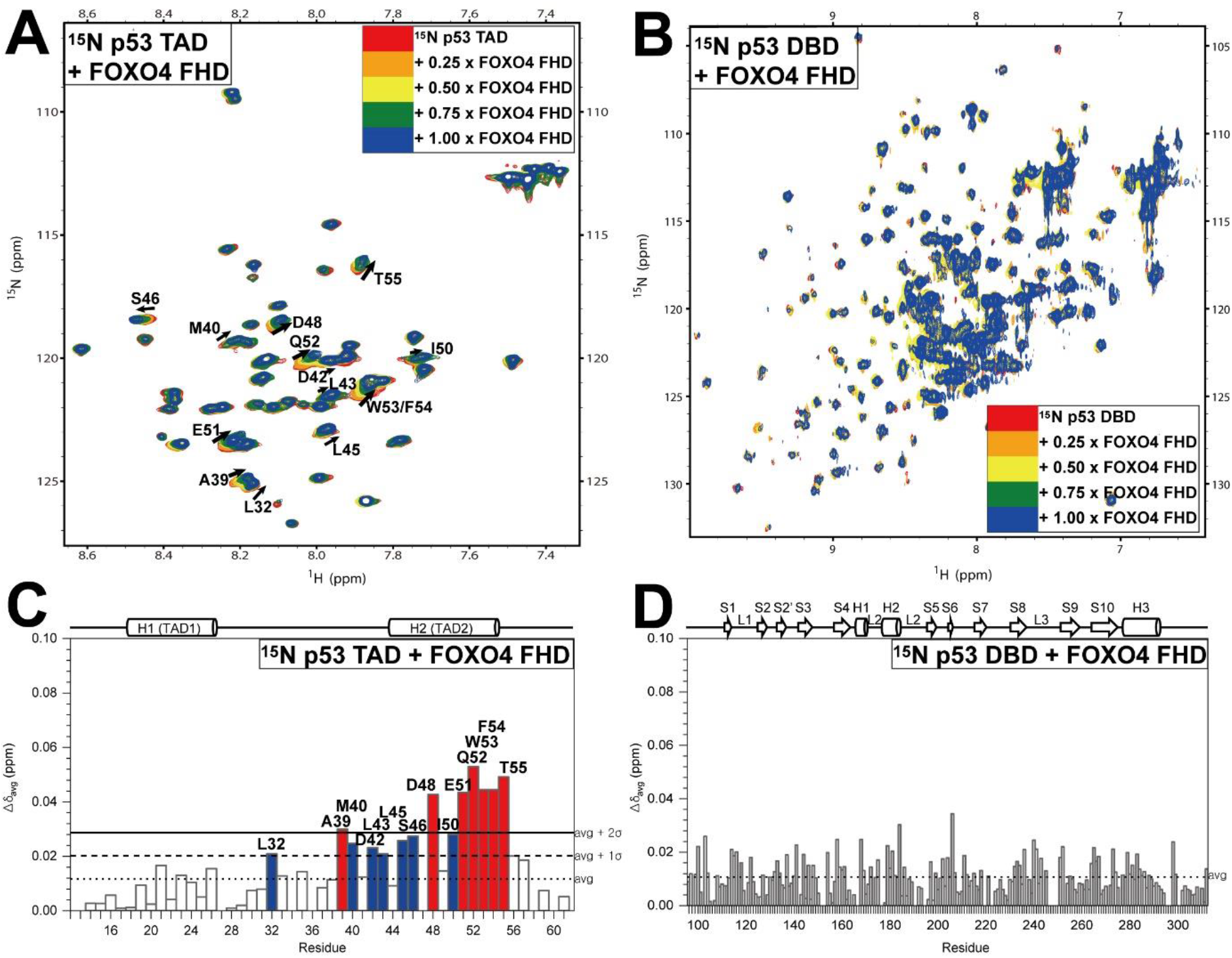
Chemical shift changes of p53 domains upon titration with FOXO4 FHD. (A) Overlaid ^1^H-^15^N HSQC spectra of 300 μM of (A) ^15^N-labeled p53 TAD and (B) ^15^N-labeled p53 DBD with increasing concentrations (0 μM, 75 μM, 150 μM, 225 μM, 300 μM) of FOXO4 FHD. (C-D) Δδ_avg_ for each (C) p53 TAD and (D) p53 DBD residue when FOXO4 FHD concentration is 300 μM. Residues that undergo a large chemical shift change are colored in red (above average + 2σ) and blue (above average + 1σ).

We additionally performed an NMR titration between ^15^N-labeled p53 DBD and FOXO4 FHD but did not find any significantly perturbed residues of p53 DBD (Figure 3B and 3D), similar to the previous experiment with ^15^N-labeled with FOXO4 FHD (Figure 2B and 2D). Our NMR data show that p53 TAD, and in particular p53 TAD2, governs the interaction with FOXO4 FHD.

### FOXO4 CR3 also participates in an interaction with p53

FOXO4 CR3 is an intrinsically disordered region, which is considered challenging to study structurally. However, we successfully observed that aliphatic amino acids of FOXO4 CR3 participate in an intramolecular interaction with FOXO4 FHD at the atomic level.[23, 49] This shows that the intrinsically disordered FOXO4 CR3 can be involved in interplay with other elements, including the intramolecular unit. Previous studies have shown that the FOXO3a CR3 interacts with intermolecular partners, and p53 DBD is one of them.[38, 39] FOXO4 and FOXO3a CR3 domains have high sequence homologies.[23, 49] Thus, we investigated whether the FOXO4 CR3 is a potential binding partner of p53 DBD. ^15^N-labeled FOXO4 CR3 was prepared and used to perform NMR titration and CSP analysis with p53 DBD. The previously reported backbone assignment of FOXO4 CR3 was applied.[23, 49] As shown in Figure 4, the CSPs of FOXO4 CR3 residues M476, I487, S488, and M491 were greater than the average + 1σ. These are mostly aliphatic amino acids, from which we can infer that FOXO4 CR3 and p53 DBD interact by hydrophobic attraction. Residues Y475, N478, L479, E480, C481, D484, I486, L490, and D492 also appeared to undergo above-average perturbations. This list is a mixture of aliphatic and acidic amino acids, suggesting that the interaction between FOXO4 TAD and p53 DBD is maintained by both hydrophobic and electrostatic attractions, similar to FOXO4 FHD-p53 TAD binding.

**Figure 4.**
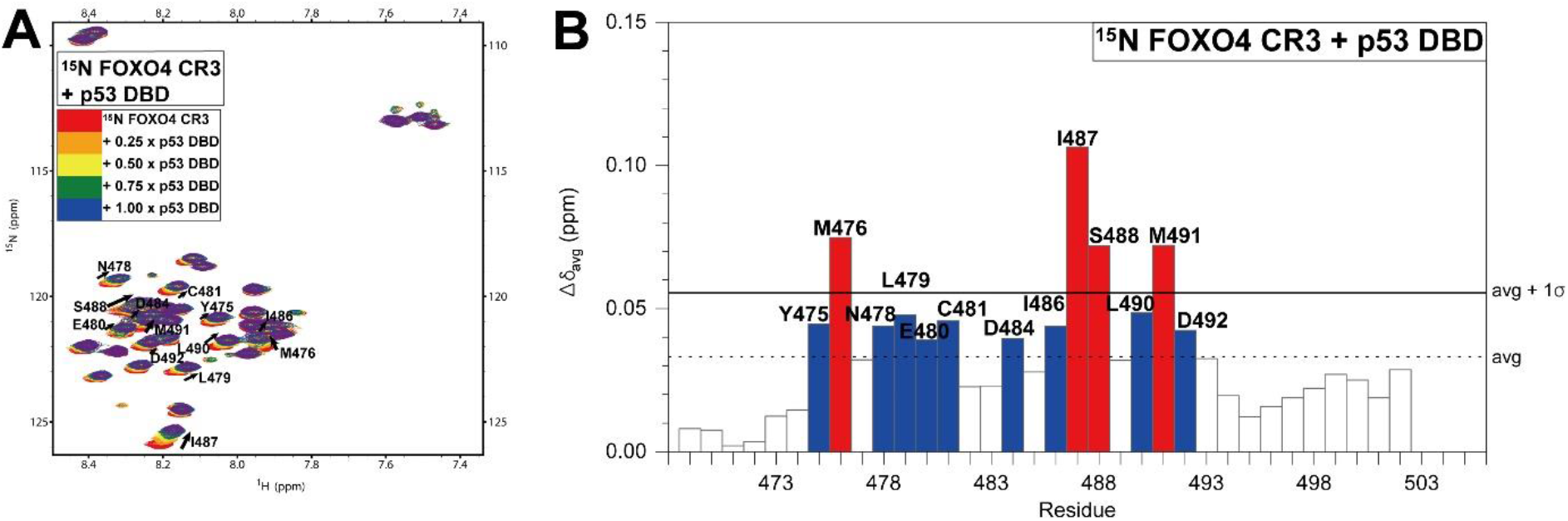
Chemical shift changes of FOXO4 CR3 upon titration with p53 DBD. (A) Overlaid ^1^H-^15^N HSQC spectra of ^15^N-labeled FOXO4 CR3 at a concentration of 300 μM with increasing concentrations (0 μM, 75 μM, 150 μM, 225 μM, 300 μM) of p53 DBD. (B) Δδ_avg_ for each FOXO4 CR3 residue when p53 DBD concentration is 300 μM. Residues that undergo a large chemical shift change are colored in red (above average + 1σ) and blue (above average).

### Binding surfaces of p53 DBD to DNA, p53 TAD and FOXO4 CR3 overlap

The main role of p53 DBD is recognition of its specific DNA sequence (5’- PuPuPuC(A/T)(T/A)GPyPyPy-3’ (Pu, purine; Py, pyrimidine)).[53] Also, the domain participates in an intramolecular PPI with p53 TAD, and intermolecular PPIs with BcL_-XL_ [54] and Hsp90.[55] Thus, p53 DBD is considered one of the vital domains for PPI of p53.

We characterized the FOXO4 CR3 binding region in p53 DBD by titrating FOXO4 CR3 into ^15^N-labeled p53 DBD. The assignment of the ^1^H-^15^N HSQC spectrum was based on previous research.[56] Y126 and S127 in β-strand S2 experienced large CSPs of more than the average + 2σ, and A138 between S2’ and S3 also had significant CSP (Figure 5A and 5B). α-Helix H3 and its surrounding residues C277, K291, K292, and G293 also underwent substantial perturbations. Several residues that changed by the average + 1σ or more were also observed, but these were spread throughout the structure. In the 3D solution structure of p53 (PDB ID: 2FEJ),[57] the positions of α-helix H3 and β-strand S2 are adjacent to each other (Figure 5C). Our CSP data suggest that p53 DBD interacts with FOXO4 CR3 through its H3 and S2 regions and neighboring surfaces. Moreover, perturbation of the basic amino acids of α-helix H3, K291, K292, and R283, indicates that electrostatic attractions are involved in the interaction between the two domains, which is consistent with the insight of the NMR titration experiment between ^15^N-labeled FOXO4 CR3 and p53 DBD (Figure 4).

**Figure 5.**
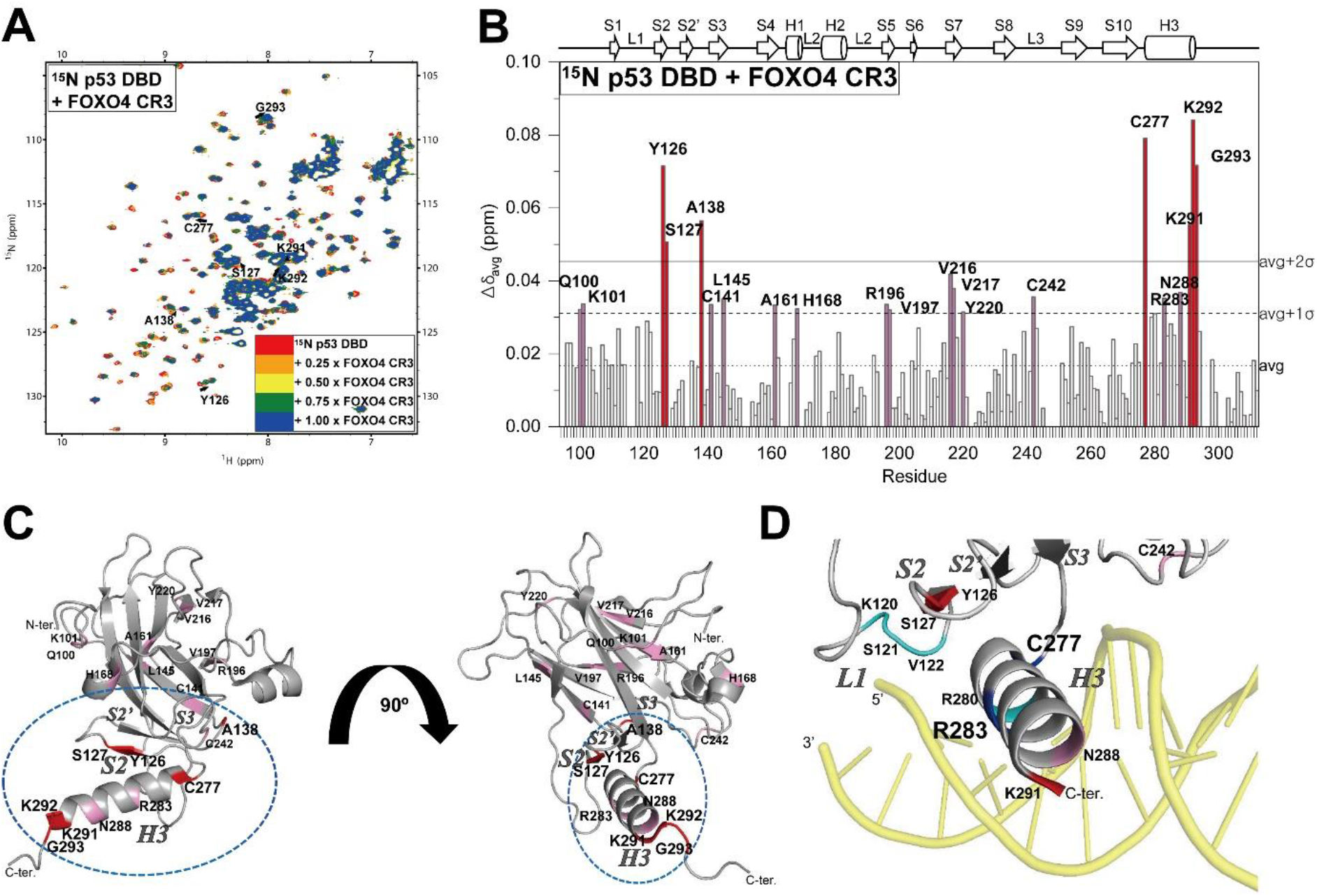
Chemical shift changes of p53 DBD upon titration with FOXO4 CR3. (A) Overlaid ^1^H-^15^N HSQC spectra of ^15^N-labeled p53 DBD at a concentration of 300 μM with increasing concentrations (0 μM, 75 μM, 150 μM, 225 μM, 300 μM) of FOXO4 CR3. (B) Δδ_avg_ for each p53 DBD residue when FOXO4 CR3 concentration is 300 μM. (C) The solution structure of p53 DBD (PDB ID: 2FEJ)[57], colored based on CSP upon addition of FOXO4 CR3. (D) The solution structure of p53 DBD with DNA (PDB ID: 3Q05).[58] Residues that undergo a large chemical shift change are colored in red (above average + 2σ) and pink (above average + 1σ). Residues that participate only in DNA binding are colored in cyan, and those involved in the binding of both DNA and FOXO4 CR3 are shown in blue.

It is noteworthy that the α-helix H3 of p53 DBD participates in the binding of FOXO4 CR3 because this α-helix also participates in the specific DNA binding of p53. In the crystal structure of p53 in complex with DNA (PDB ID: 3Q05) (Figure 5D)[58], K120 and S121 in loop L1 of the DBD, and C277, R280, and R283 in α-helix H3 compose the DNA binding surface. Among these, C277 and R283 showed large CSPs with FOXO4 CR3. In addition, it has been shown that p53 TAD interacts with the α-helix H3 region of p53 DBD and plays a role in regulating target DNA recognition of p53.[52, 59] In particular, residues such as K291 and G293 of p53 DBD that were significantly perturbed by p53 TAD were also changed by FOXO4 CR3.[59] Overall, the overlapping of several residues of p53 DBD α-helix H3 implies that common surfaces are shared for DNA recognition, intramolecular p53 TAD binding, and intermolecular FOXO4 CR3 interaction.

### The interaction between FOXO4 CR3 and p53 DBD is mediated over a wide surface

In the above experiments, the largest CSPs in FOXO4 CR3 induced by p53 DBD were limited to four residues, M476, I487, S488, and M491 (Figure 4B). This differs from the experiment with ^15^N-labeled p53 TAD and FOXO4 FHD, where large CSPs were concentrated in one α-helix (TAD2) (Figure 3C). We hypothesized that because of the intrinsically disordered nature of FOXO4 CR3, the binding surface might not be highly refined.

In order to evaluate the role of the four most perturbed residues for the binding of p53 DBD, we introduced alanine mutations for CR3 residues M476, I487, S488, and M491. We performed NMR titration experiments with ^15^N-labeled p53 DBD (Figure 6). As shown in the CSP analysis of ^15^N-labeled p53 DBD and the FOXO4 CR3 mutants, the perturbation of residues belonging to β-strand S2 and α-helix H3, which were observed as important regions for binding to wild-type FOXO4 CR3, was relatively well maintained in mutants I487A, S488A, and M491A (Figure 6B, 6C, and 6D). Although CSPs of S127 and A138 located in β-strand S2 and its surrounding residues decreased in magnitude, the aromatic amino acid Y126 still maintained a CSP above the average + 2σ of the experiment with wild-type CR3 protein. CSPs of C277, K291, K292, and G293, which are important α-helix H3 residues in the FOXO4 CR3 binding, also maintained high values.

**Figure 6.**
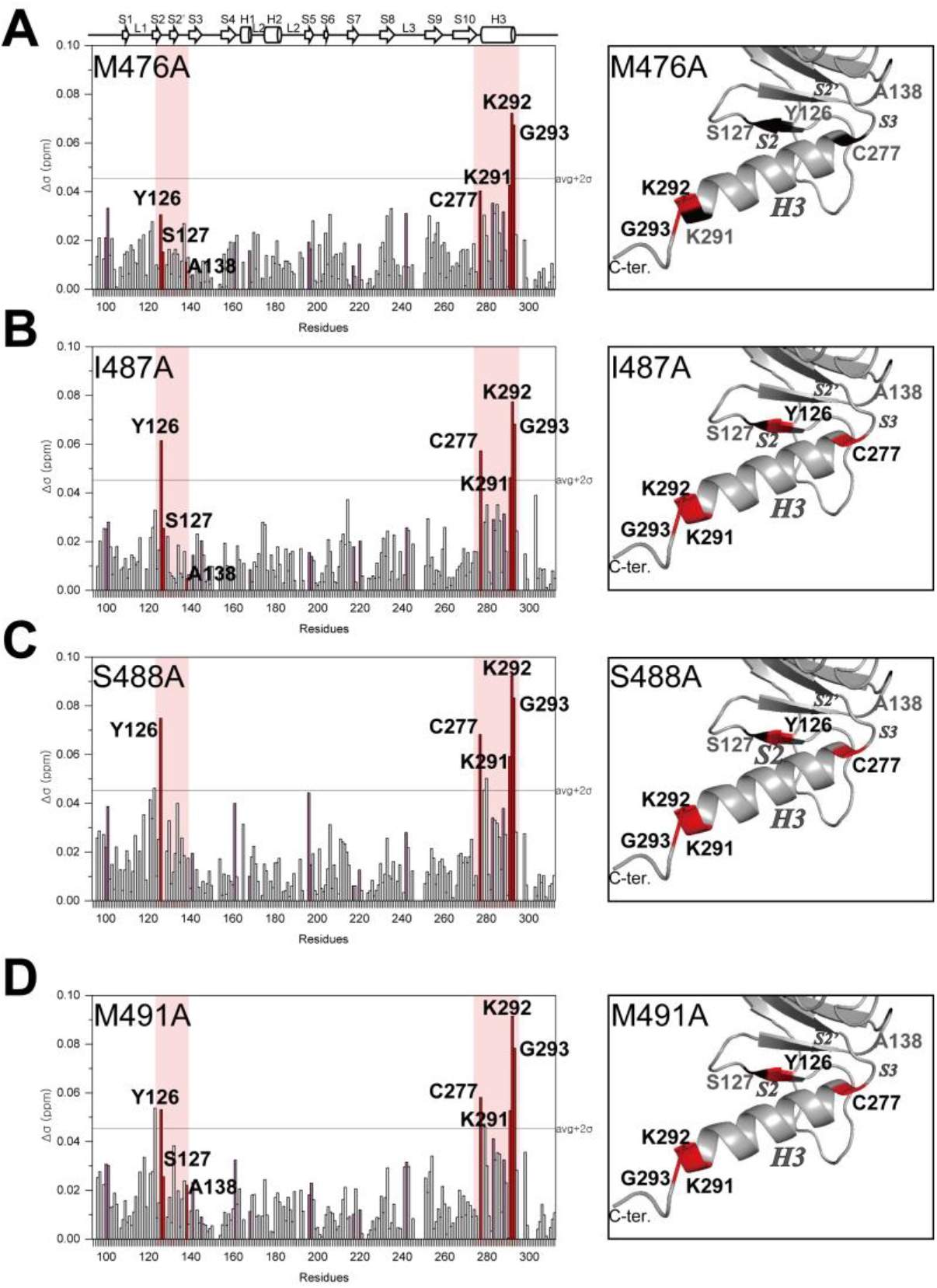
Chemical shift changes of p53 DBD after titration with FOXO4 CR3 alanine mutants. Δδ_avg_ for each ^15^N-labeled p53 DBD at a concentration of 300 μM when concentration of FOXO4 CR3 (A) M476A, (B) I487A, (C) S488A, and (D) M491A are present at 300 μM (left column). Residues that undergo a large chemical shift change in ^15^N-labeled p53 DBD and FOXO4 CR3 titration are colored in red (above average + 2σ) and pink (above average + 1σ). Vertical pink bands indicate the regions between β-strands S2 and S3, and α-helix H3 of p53 DBD, which showed large perturbations in wild-type FOXO4 CR3 titration. The solution structure of p53 DBD (PDB ID: 2FEJ)[57], colored based on CSP upon addition of the FOXO4 CR3 alanine mutants (right column). Residues that undergo significant changes in both CR3 and its mutants are colored in red, and residues that undergo large perturbation in CR3 but are reduced by mutation are colored in black.

On the other hand, the FOXO4 CR3 M476A mutant showed a different tendency than the others. In Figure 6A, the CSP of the key residues of β-strand S2 was significantly reduced. The perturbation of Y126 was markedly reduced. In addition, the CSP of residue C277, a residue structurally adjacent to β-strand S2, was also relatively small compared to the other three cases. This data indicates that M476 of FOXO4 CR3 is an important residue for the binding of p53 DBD β-strand S2. However, as the perturbation of residues belonging to α-helix H3 is relatively maintained, this mutation does not seem to abolish the binding between FOXO4 CR3 and p53 DBD completely. Based on these results, we suggest that the FOXO4 CR3-p53 DBD interface is distributed across several residues in CR3, such that the interaction is maintained if one of them is missing. Thus, it can be inferred that the flexible FOXO4 CR3, without a distinct secondary structure, can provide various contact points over a wide surface during the binding with p53 DBD.

### Thermodynamic analysis of FOXO4-p53 dual binding surface

We performed ITC experiments to quantify the thermodynamic profiles of the binding between domains that contribute to the formation of the FOXO4-p53 complex.

As shown in Figure 7A, the binding between FOXO4 FHD and p53 TAD has a dissociation equilibrium constant (*K*_d_) of 2.19 ± 0.06 μM. The ΔG and ΔH values for this reaction were calculated as −7.15 ± 0.46 kcal/mol and 12.29 ± 2.06 kcal/mol, respectively (Figure 7B). This means that the binding takes place in the form of a spontaneous endothermic reaction. Also, the value of stoichiometry (n) was determined to be 1.04, suggesting that they participate in the binding at a 1:1 ratio (Figure 7A). The positive change in entropy (ΔS) observed appears to be due to the hydrophobic attraction between the two domains causing the scatter of water molecules on the binding surface. This suggestion is supported by the NMR data, in which the perturbation of several hydrophobic amino acids was observed between FOXO FHD and p53 TAD. The signs of ΔH, ΔS, and ΔG are the same as those of the intramolecular interaction of FOXO4 (FOXO4 FHD and CR3).[23, 49] This implies that the binding of FOXO4 FHD to p53 TAD occurs with a mechanism similar to contact with FOXO4 CR3.

**Figure 7.**
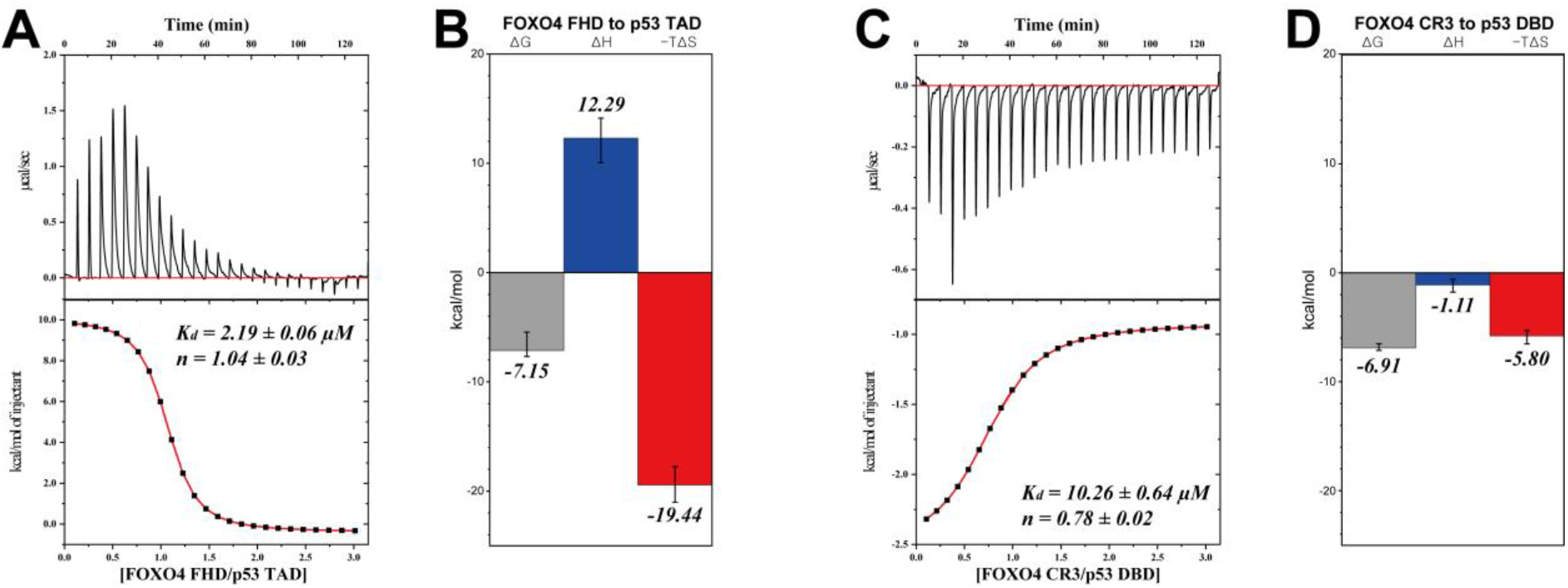
Thermodynamic analysis of FOXO4-p53 domain interactions by ITC. Raw data of heat plot (upper panel) and integration plot of data (lower panel) when (A) FOXO4 FHD was injected into p53 TAD and (C) FOXO4 CR3 was injected into p53 DBD. Binding affinity (*K*_d_) and stoichiometry (n) between the two molecules in each sample are presented. Three thermodynamic parameters (ΔH, enthalpy change; ΔG, Gibbs energy of association; −TΔS, entropy change times temperature) of (B) FOXO4 FHD-p53 TAD interaction and (D) FOXO4 CR3-p53 DBD interaction obtained from the curve.

FOXO4 CR3 and p53 DBD interact with different thermodynamic characteristics. The *K*_d_ was measured to be 10.26 ± 0.64 μM, and ΔG and ΔH were −6.91 ± 0.35 kcal/mol and −1.11 ± 0.61 kcal/mol, which is assumed to be a spontaneous exothermic reaction. The value of n was 0.781, which is a little less than 1, but it can be assumed that the two domains interact in a 1:1 ratio. The ΔS was positive, the same as in the FOXO4 FHD-p53 TAD experiment, but the magnitude of the change was smaller. The FOXO4 FHD-p53 TAD interaction has a *K*_d_ value about 4.7 times smaller than the FOXO4 CR3-p53 DBD interaction. This comparison suggests that when FOXO4 and p53 are in close proximity, the binding of FOXO4 FHD-p53 TAD takes precedence over the FOXO4 CR3-p53 DBD binding.

## Discussion

FOXO4 and p53 play a pivotal role in cellular senescence/apoptosis.[11, 26] In senescent cells, FOXO4 is elevated and contributes to maintaining cell viability.[11] As senescence progresses, FOXO4 and p53 co-localize and share roles such as inducing transcription of p21.[11, 34] Thus, the interaction between FOXO4 and p53 is expected to play a direct role in cellular senescence. In fact, a peptide-based inhibitor that mimics a part of FOXO4 FHD induced p53-dependent apoptosis through binding to p53.[11] Therefore, to understand the mechanism of cell lifespan and to control it appropriately, information on the molecular level of FOXO4-p53 binding is required. In this study, we expanded the structural understanding of the interaction between FOXO4 and p53.

In previous studies, the FOXO4 FHD binding region of p53 was briefly determined as amino acids 1-312[11], and the intramolecular interaction between p53 TAD and DBD was demonstrated.[52] Here, we specified that the p53 TAD2 is a crucial region for the interaction with FOXO4 FHD (Figure 3C). We found that p53 TAD binds to the pocket-like space between the N-terminal loop of FOXO4 FHD and α-helices H1, H3, and H4 through NMR titration experiments (Figure 2E). We previously have shown that the FOXO4 CR3 can bind intramolecularly to a surface formed by the N-terminal loop, α-helices H1, and H3, β-strand S2, and wing W1 of FOXO4 FHD.[23, 49] This suggests that most of the p53 TAD binding region of FOXO4 FHD is contained in the FOXO4 CR3 binding region. The sites even have large CSP values in common for Q104 of the N-terminal loop, S105 of α-helix H1, and L160 of α-helix H3. While FOXO4 CR3 and p53 TAD2 partially share the binding surfaces of FOXO4 FHD, the thermodynamic parameters measured by ITC experiments using isolated constructs showed that the FOXO4 CR3-FHD interaction has a higher binding affinity *in vitro* than the p53 TAD-FOXO4 FHD complex. Table 1 shows the *K*_d_ and ΔG values of both interactions. These differences imply that the intramolecular binding between the two domains of FOXO4, FHD and CR3, is more substantial and forms a thermodynamically stable complex. However, the higher equilibrium binding affinity of FOXO4’s intramolecular interaction does not rule out the possibility of FOXO4 FHD-p53 TAD binding. From a kinetic perspective, FOXO4 FHD associates with and dissociates from FOXO4 CR3, and in its free state, FOXO4 FHD has a chance to interact with p53 TAD. With the enhanced expression level of FOXO4 in senescent cells [11], the FOXO4 FHD-p53 TAD interaction could be biologically significant to inhibit the release of p53 out of the nucleus and eventually to maintain cellular senescence.

**Table 1.**
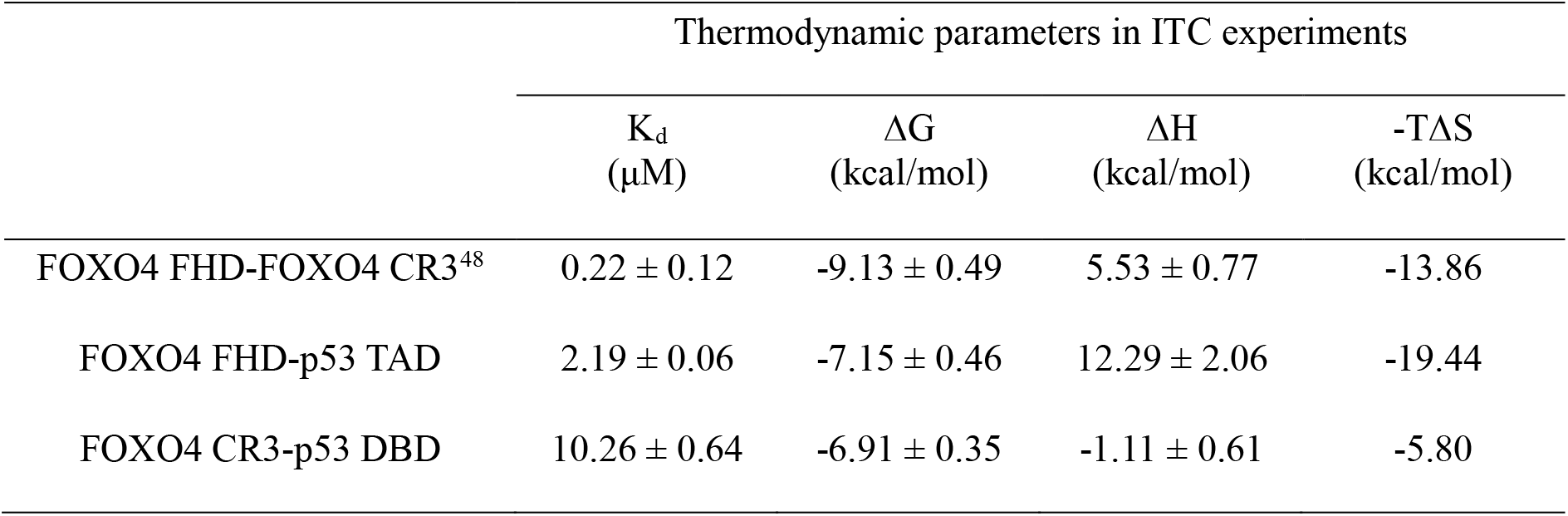
Thermodynamic parameters of FOXO4 intramolecular interaction and FOXO4-p53 intermolecular interaction.

We newly identified an additional point of contact between FOXO4 and p53 in this study. The binding of FOXO4 CR3 and p53 DBD has significance in that it can provide the second junction that can enhance the FOXO4-p53 interaction in addition to the primary binding between FOXO4 FHD and p53 TAD. NMR titrations through reciprocal ^15^N labeling showed that four FOXO4 CR3 amino acids (M476, I487, S488, and M491) are required to interact with the p53 DBD α-helix H3 and its surrounding residues (Figures 4 and 5). α-Helix H3 of p53 DBD participates in both DNA binding (Figure 5D)[58] and the interaction with p53 TAD[52], and this helix is in one of the pivotal regions for the various other molecular interactions of p53. While FOXO4 CR3 and p53 TAD contact similar surfaces on p53 DBD, the *K*_d_ of FOXO4 CR3-p53 DBD shows that the binding affinity is relatively low (Table 1, Figure 7B). Taken together, the two intermolecular interactions between FOXO4 and p53 are weaker compared to each molecule’s intramolecular interaction. However, in conditions of FOXO4 overcrowding, such as in senescent cells, an additional point of contact for the FOXO4-p53 interaction may be sufficient to overcome the strength of the intramolecular interactions. It has been recognized that weak and transient PPIs with micromolar to millimolar range binding affinity can play an important role in cell signaling and regulatory and response mechanisms by modulating the bound and unbound state populations between two molecules.[60–62] In this regard, we suggest that binding between FOXO4 CR3 and p53 DBD also can be a factor responsible for the maintenance of senescent cells along with the FOXO4 FHD-p53 TAD interaction.

As TFs, p53 and FOXO4 recognize their cognate DNA elements, and intramolecular interactions between p53 DBD-TAD and FOXO4 FHD-CR3 affect their sequence-specific DNA binding properties.[23, 52] It is known that both FOXO4 CR3 and p53 TAD are released from intramolecular interactions upon target DNA binding of FHD/DBD.[23, 52] In this aspect, DNA binding of either FOXO4 or p53 could expose transactivation domain which mediates intermolecular interaction with the other. Previous studies showed that the cooperative binding of several TFs in the DNA transcription process is necessary.[63, 64] The cooperation between FOXO4 and p53 seems essential for the transcription of p21, which accelerates cellular senescence. Target DNA elements for FOXO4 and two p53s are adjacent to one another within the promoter site of *CDKN1A*, the gene encoding p21.[11, 31] Also, it has been shown that a peptide inhibitor of FOXO4-p53 binding significantly reduced the expression level of p21.[11] These suggest that the FOXO4-p53 co-localization on the promoter site affects the transcriptional activity of p21, a prerequisite for cellular senescence that occurs in response to DNA damage (Figure 8A). From this point of view, we suggest that the dual binding surface between FOXO4 and p53 has a potentially synergistic effect for locating two TFs on their respective target DNA sequences in the preparation stage of senescent cell transcription through the following three models (Figure 8B): (a) When p53 is bound to the p21 promoter, p53 DBD is in contact with DNA, so p53 TAD is freely exposed in the nucleus and recruits FOXO4 to the p21 promoter through interaction with FOXO4 FHD. Afterward, FOXO4 searches for its target sequence in the p21 promoter through rearrangement. (b) When FOXO4 FHD is bound to the p21 promoter, FOXO4 CR3 exists in the free state and has an opportunity to attach to p53 DBD. In turn, p53 DBD is transferred to the p53 binding site of the p21 promoter. (c) When FOXO4 and p53 exist in a complex and are close to the p21 promoter region, the FOXO4 CR3-p53 DBD interaction of relatively low binding affinity is removed first, and the free p53 DBD binds to the target DNA sequence. Subsequently, the binding between FOXO4 FHD and p53 TAD is released, allowing FOXO4 FHD to recognize its promoter site. As a result, freely exposed p53 TAD and FOXO4 CR3 can bind to transcription co-activators such as CBP/p300 and initiate transcription of p21. We hypothesize that this dual binding system between these two TFs can make initiation of p21 transcription more efficient because it provides ways to recruit the second TF when the first one is already at the promoter region. Nevertheless, details of the structural and biophysical characteristics of tertiary complexes composed of FOXO4, p53, and DNA promoters are still unknown, so future research on the FOXO4-p53-DNA binding mechanism will be needed.

**Figure 8.**
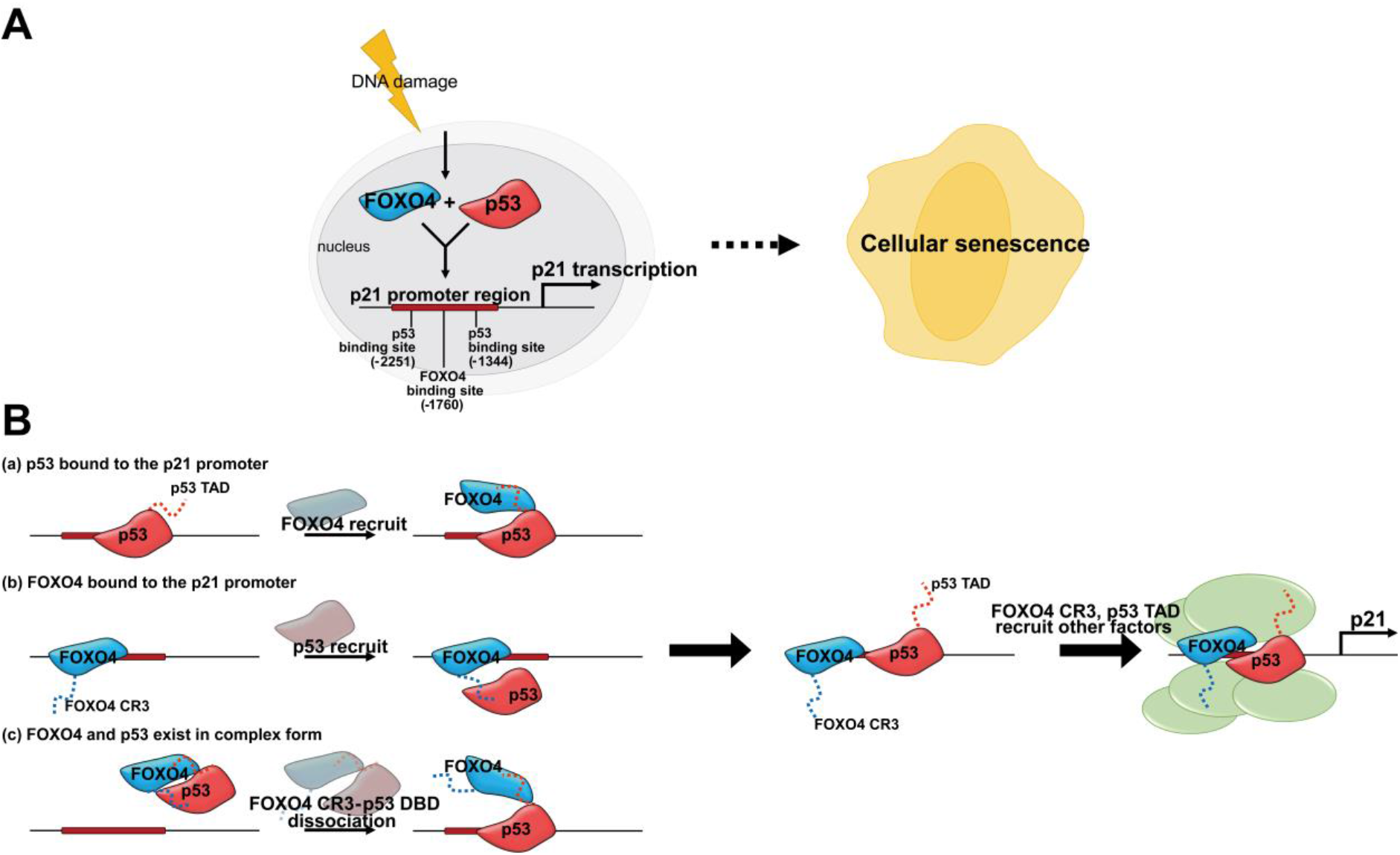
(A) FOXO4 and p53 regulate cellular senescence by recognizing the promoter region of p21. (B)Model for cooperative transcription initiation of FOXO4 and p53.

As mentioned above, the development of an inhibitor that interferes with the FOXO4-p53 interaction has recently been reported. The FOXO4-DRI peptide, a mimetic of the FOXO4 FHD N-terminus, defended cells from senescence.[11] Our FOXO4 FHD-p53 TAD NMR titration results (Figure 2C and 3C) suggest that FOXO4-DRI peptide exerts its inhibitory effect through binding to p53 TAD2. In other words, this inhibitor appears to regulate p53 activation in senescent cells by interrupting contact with co-activators of p53 instead of with DNA binding. It will be interesting to conduct studies that introduce and apply this inhibitor to deal with various p53-related diseases. On the other hand, we found in this study that the interaction of FOXO4 and p53 involves a dual interface. This implies that the binding of FOXO4 CR3-p53 DBD can be targeted along with FOXO4 FHD-p53 TAD to inhibit their interaction. We observed that four residues of FOXO4 CR3 are important for the interaction with p53 (Figure 4), but it seems that there is no dominant involvement of a specific residue because this interaction is expected to be across a wide surface (Figure 6). This implies that inhibitors targeting the FOXO4 CR3-p53 DBD binding are difficult to achieve with small molecules,[65] and that peptide mimics would be better candidates. Therefore, appropriate modification of the relatively nonessential residues of FOXO4 CR3 except for the four amino acids that were sensitive to the presence of p53 DBD is recommended for the purpose of drug development that interferes with PPIs of p53. α-Helix induction through leucine substitution is a good example.[66] Moreover, introducing an additional FOXO4 CR3 sequence, which is important for p53 binding, next to the FOXO4 DRI peptide is expected to be a reasonable candidate for making more precise and specific p53 inhibitors.

## Conclusions

Cellular senescence is a process that protects against unexpected oncogenic stress in somatic cells. However, the accumulation of senescent cells causes aging and diseases such as cancer. Understanding protein interactions involved in cellular senescence could provide a clue to elucidating its cause and overcoming disease through appropriate targeting. Through the study of the FOXO4-p53 axis, we observed that the FOXO4 and p53 interaction has dual binding surfaces, providing their DNA binding and transactivation domains as contact points. In addition, a model is presented for how the interaction of two TFs cooperates with the transcription initiation of a downstream gene. This study could provide essential structural information for drug development to overcome senescence-associated diseases.

## Methods

### Sample preparation

The full-length FOXO4 and p53 genes were acquired from Addgene. FOXO4 FHD (95-195) and p53 DBD (94-312) were subcloned into the pET His6 TEV LIC cloning vector (2B-T) (a gift from Scott Gradia, Addgene plasmid #29666) and transformed into *E. coli* BL21 (DE3) cells. FOXO4 CR3 (469-505) and p53 TAD (13-61) were subcloned into the pET His6 GST TEV LIC cloning vector (2G-T) (a gift from Scott Gradia, Addgene plasmid #29707) and transformed into *E. coli* BL21 (DE3) pLysS cells. All cells were grown at 37 °C to an OD_600_ of 0.7 in Luria-Bertani (LB) medium, and 0.5 mM of isopropyl-1-thio-β-D-galactopyranoside was added. Cells were cultured overnight (20 h) at 18 °C after additional induction. Overexpressed proteins were purified on a Ni-NTA column (GE Healthcare) followed by gel filtration chromatography using Hi-Load 16/600 superdex 200 pg and 75 pg (GE Healthcare) on AKTA pure and AKTA prime systems. The composition of the buffer solution was 20 mM HEPES (pH 7.0), 50 mM NaCl, 0.5 mM DTT. Glutathione-S-transferase (GST) tagged recombinant protein samples were cleaved by Tobacco Etch Virus (TEV) protease. The four FOXO4 CR3 variants (M476A, I487A, S488A, and M491A) were generated from the *FOXO4* gene by a site-directed mutagenesis strategy. ^15^N-labeled protein samples for ^1^H-^15^N HSQC experiments were obtained by culturing cells in M9 minimal media containing ^15^N-labeled NH_4_Cl. They had the same overexpression and purification process as cells cultured in LB media.

### Nuclear magnetic resonance experiments

Bruker 900 MHz (KBSI, Ochang) and 600 MHz (GIST, Gwangju) NMR spectrometers with cryogenic probes were used to conduct NMR experiments. The buffer solution conditions for all experiments were set to 20 mM NaPi (pH 7.0), 50 mM NaCl, 2 mM DTT, and the temperature was set to 25 °C. Data processing and analysis were performed using Topspin (Bruker) and NMRFAM-Sparky software programs.[67] The amide nitrogens and protons of FOXO4 FHD,[48] FOXO4 CR3,[23, 49] and p53 DBD[56] have been assigned previously. The backbone assignments of p53 TAD were obtained by performing ^1^H-^15^N HSQC, HNCO, HNCACO, HNCACB, and CBCACONH experiments at a concentration of 1.0 mM. ^1^H-^15^N HSQC titration experiments were performed by adding 0 μM, 75 μM, 150 μM, 225 μM, and 300 μM unlabeled protein to ^15^N-labeled samples of 300 μM. Average CSP values (*Δδ_avg_*) were calculated using the following equation:

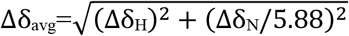

Δδ_H_ and Δδ_N_ are the chemical shift changes of the amide proton and nitrogen, respectively.

### Isothermal titration calorimetry analysis

ITC experiments were performed in 20 mM HEPES (pH 7.0), 50 mM NaCl, 0.5 mM DTT buffer solution using a Nano-ITC SV instrument (GIST, Gwangju). Titration was performed by adding 10 μL of 1000 μM protein via syringe injection into 100 μM protein in the cell (1000 μL) 25 times. The temperature in all experiments was 25 °C; the first injection (10 μL) was not used in the analysis. The time interval between injections was 300-350 sec, and the stirring speed was 350 rpm. Error values of the result were derived from the results of 3 replicate experiments. The thermodynamic data (stoichiometry (n), dissociation constant (*K*_d_), enthalpy change (ΔH), and entropy change (ΔS)) were analyzed using the ‘Independent model’ fitting of the NanoAnalyze software (TA Instruments). The Gibbs free energy change (ΔG) was calculated by the following equation:

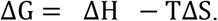

## Abbreviations

FOXO4: forkhead box O 4
FHD: forkhead domain
TAD: transactivation domain
DBD: DNA binding domain
NMR: nuclear magnetic resonance
HSQC: heteronuclear single quantum coherence
CSP: chemical shift perturbation
ITC: isothermal titration calorimetry
PPI: protein-protein interaction
TF: transcription factor

## Acknowledgments

We thank the high-field NMR facility at the Korea Basic Science Institute (KBSI, Ochang) and GIST Central Research Facilities (GIST, Gwangju) for allowing us to use their NMR spectrometers. This work was supported by the National Research Foundation of Korea [grant 2018R1A2B6004388 and 2021R1A2C1004669 to C.-J.P], which is funded by the Korean government (MSIT); by a GIST Research Institute grant funded by GIST in 2021; and by the KBSI under the R&D program [Project No.C140440] supervised by the Ministry of Science and ICT, Korea.

## Author Contributions

J.K., D.A., and C.-J.P. designed the study; J.K. and D.A. prepared samples; J.K., D.A., and C.-J.P. planned and performed experiments; J.K., D.A., and C.-J.P. analyzed the data; J.K., D.A., and C.-J.P. wrote the paper. All authors have given approval to the final version of the manuscript.

## References

1. Myatt, S. S. & Lam, E. W. (2007) The emerging roles of forkhead box (Fox) proteins in cancer, Nature reviews Cancer. 7, 847–59.

2. Carter, M. E. & Brunet, A. (2007) FOXO transcription factors, Current biology : CB. 17, R113–4.

3. Accili, D. & Arden, K. C. (2004) FoxOs at the crossroads of cellular metabolism, differentiation, and transformation, Cell. 117, 421–6.

4. Barthel, A., Schmoll, D. & Unterman, T. G. (2005) FoxO proteins in insulin action and metabolism, Trends in endocrinology and metabolism: TEM. 16, 183–9.

5. Birkenkamp, K. U. & Coffer, P. J. (2003) Regulation of cell survival and proliferation by the FOXO (Forkhead box, class O) subfamily of Forkhead transcription factors, Biochemical Society transactions. 31, 292–7.

6. Essers, M. A., Weijzen, S., de Vries-Smits, A. M., Saarloos, I., de Ruiter, N. D., Bos, J. L. & Burgering, B. M. (2004) FOXO transcription factor activation by oxidative stress mediated by the small GTPase Ral and JNK, The EMBO journal. 23, 4802–12.

7. de Keizer, P. L., Packer, L. M., Szypowska, A. A., Riedl-Polderman, P. E., van den Broek, N. J., de Bruin, A., Dansen, T. B., Marais, R., Brenkman, A. B. & Burgering, B. M. (2010) Activation of forkhead box O transcription factors by oncogenic BRAF promotes p21cip1-dependent senescence, Cancer research. 70, 8526–36.

8. van den Berg, M. C., van Gogh, I. J., Smits, A. M., van Triest, M., Dansen, T. B., Visscher, M., Polderman, P. E., Vliem, M. J., Rehmann, H. & Burgering, B. M. (2013) The small GTPase RALA controls c-Jun N-terminal kinase-mediated FOXO activation by regulation of a JIP1 scaffold complex, The Journal of biological chemistry. 288, 21729–41.

9. Nagaich, A. K., Zhurkin, V. B., Durell, S. R., Jernigan, R. L., Appella, E. & Harrington, R. E. (1999) p53-induced DNA bending and twisting: p53 tetramer binds on the outer side of a DNA loop and increases DNA twisting, Proceedings of the National Academy of Sciences of the United States of America. 96, 1875–80.

10. Brownawell, A. M., Kops, G. J., Macara, I. G. & Burgering, B. M. (2001) Inhibition of nuclear import by protein kinase B (Akt) regulates the subcellular distribution and activity of the forkhead transcription factor AFX, Molecular and cellular biology. 21, 3534–46.

11. Baar, M. P., Brandt, R. M. C., Putavet, D. A., Klein, J. D. D., Derks, K. W. J., Bourgeois, B. R. M., Stryeck, S., Rijksen, Y., van Willigenburg, H., Feijtel, D. A., van der Pluijm, I., Essers, J., van Cappellen, W. A., van, I. W. F., Houtsmuller, A. B., Pothof, J., de Bruin, R. W. F., Madl, T., Hoeijmakers, J. H. J., Campisi, J. & de Keizer, P. L. J. (2017) Targeted Apoptosis of Senescent Cells Restores Tissue Homeostasis in Response to Chemotoxicity and Aging, Cell. 169, 132–147.e16.

12. Coppé, J. P., Patil, C. K., Rodier, F., Sun, Y., Muñoz, D. P., Goldstein, J., Nelson, P. S., Desprez, P. Y. & Campisi, J. (2008) Senescence-associated secretory phenotypes reveal cell-nonautonomous functions of oncogenic RAS and the p53 tumor suppressor, PLoS biology. 6, 2853–68.

13. Rodier, F., Coppé, J. P., Patil, C. K., Hoeijmakers, W. A., Muñoz, D. P., Raza, S. R., Freund, A., Campeau, E., Davalos, A. R. & Campisi, J. (2009) Persistent DNA damage signalling triggers senescence-associated inflammatory cytokine secretion, Nature cell biology. 11, 973–9.

14. McHugh, D. & Gil, J. (2018) Senescence and aging: Causes, consequences, and therapeutic avenues, The Journal of cell biology. 217, 65–77.

15. Rosso, A., Balsamo, A., Gambino, R., Dentelli, P., Falcioni, R., Cassader, M., Pegoraro, L., Pagano, G. & Brizzi, M. F. (2006) p53 Mediates the accelerated onset of senescence of endothelial progenitor cells in diabetes, The Journal of biological chemistry. 281, 4339–47.

16. Jeon, O. H., Kim, C., Laberge, R. M., Demaria, M., Rathod, S., Vasserot, A. P., Chung, J. W., Kim, D. H., Poon, Y., David, N., Baker, D. J., van Deursen, J. M., Campisi, J. & Elisseeff, J. H. (2017) Local clearance of senescent cells attenuates the development of post-traumatic osteoarthritis and creates a pro-regenerative environment, Nature medicine. 23, 775–781.

17. Gorgoulis, V. G., Pratsinis, H., Zacharatos, P., Demoliou, C., Sigala, F., Asimacopoulos, P. J., Papavassiliou, A. G. & Kletsas, D. (2005) p53-dependent ICAM-1 overexpression in senescent human cells identified in atherosclerotic lesions, Laboratory investigation; a journal of technical methods and pathology. 85, 502–11.

18. Querfurth, H. W. & LaFerla, F. M. (2010) Alzheimer’s disease, The New England journal of medicine. 362, 329–44.

19. Bansal, R. & Nikiforov, M. A. (2010) Pathways of oncogene-induced senescence in human melanocytic cells, Cell cycle (Georgetown, Tex). 9, 2782–8.

20. Klotz, L. O., Sánchez-Ramos, C., Prieto-Arroyo, I., Urbánek, P., Steinbrenner, H. & Monsalve, M. (2015) Redox regulation of FoxO transcription factors, Redox biology. 6, 51–72.

21. Biggs, W. H., 3rd, Cavenee, W. K. & Arden, K. C. (2001) Identification and characterization of members of the FKHR (FOX O) subclass of winged-helix transcription factors in the mouse, Mammalian genome : official journal of the International Mammalian Genome Society. 12, 416–25.

22. Furuyama, T., Nakazawa, T., Nakano, I. & Mori, N. (2000) Identification of the differential distribution patterns of mRNAs and consensus binding sequences for mouse DAF-16 homologues, The Biochemical journal. 349, 629–34.

23. Kim, J., Ahn, D., Park, CJ. (2021) FOXO4 transactivation domain interaction with forkhead DNA binding domain and effect on selective DNA recognition for transcription initiation, Journal of molecular biology. in press.

24. Hartlmüller, C., Spreitzer, E., Göbl, C., Falsone, F. & Madl, T. (2019) NMR characterization of solvent accessibility and transient structure in intrinsically disordered proteins, Journal of biomolecular NMR. 73, 305–317.

25. Shtutman, M., Chang, B. D., Schools, G. P. & Broude, E. V. (2017) Cellular Model of p21-Induced Senescence, Methods in molecular biology (Clifton, NJ). 1534, 31–39.

26. Kruiswijk, F., Labuschagne, C. F. & Vousden, K. H. (2015) p53 in survival, death and metabolic health: a lifeguard with a licence to kill, Nature reviews Molecular cell biology. 16, 393–405.

27. Liu, G., Parant, J. M., Lang, G., Chau, P., Chavez-Reyes, A., El-Naggar, A. K., Multani, A., Chang, S. & Lozano, G. (2004) Chromosome stability, in the absence of apoptosis, is critical for suppression of tumorigenesis in Trp53 mutant mice, Nature genetics. 36, 63–8.

28. Rowan, S., Ludwig, R. L., Haupt, Y., Bates, S., Lu, X., Oren, M. & Vousden, K. H. (1996) Specific loss of apoptotic but not cell-cycle arrest function in a human tumor derived p53 mutant, The EMBO journal. 15, 827–38.

29. Toledo, F., Krummel, K. A., Lee, C. J., Liu, C. W., Rodewald, L. W., Tang, M. & Wahl, G. M. (2006) A mouse p53 mutant lacking the proline-rich domain rescues Mdm4 deficiency and provides insight into the Mdm2-Mdm4-p53 regulatory network, Cancer cell. 9, 273–85.

30. Vousden, K. H. & Prives, C. (2009) Blinded by the Light: The Growing Complexity of p53, Cell. 137, 413–31.

31. Bourgeois, B. & Madl, T. (2018) Regulation of cellular senescence via the FOXO4-p53 axis, FEBS letters. 592, 2083–2097.

32. Tran, H., Brunet, A., Grenier, J. M., Datta, S. R., Fornace, A. J., Jr., DiStefano, P. S., Chiang, L. W. & Greenberg, M. E. (2002) DNA repair pathway stimulated by the forkhead transcription factor FOXO3a through the Gadd45 protein, Science (New York, NY). 296, 530–4.

33. Seoane, J., Le, H. V., Shen, L., Anderson, S. A. & Massagué, J. (2004) Integration of Smad and forkhead pathways in the control of neuroepithelial and glioblastoma cell proliferation, Cell. 117, 211–23.

34. Miyauchi, H., Minamino, T., Tateno, K., Kunieda, T., Toko, H. & Komuro, I. (2004) Akt negatively regulates the in vitro lifespan of human endothelial cells via a p53/p21-dependent pathway, The EMBO journal. 23, 212–20.

35. Niculescu, A. B., 3rd, Chen, X., Smeets, M., Hengst, L., Prives, C. & Reed, S. I. (1998) Effects of p21(Cip1/Waf1) at both the G1/S and the G2/M cell cycle transitions: pRb is a critical determinant in blocking DNA replication and in preventing endoreduplication, Molecular and cellular biology. 18, 629–43.

36. Chang, B. D., Watanabe, K., Broude, E. V., Fang, J., Poole, J. C., Kalinichenko, T. V. & Roninson, I. B. (2000) Effects of p21Waf1/Cip1/Sdi1 on cellular gene expression: implications for carcinogenesis, senescence, and age-related diseases, Proceedings of the National Academy of Sciences of the United States of America. 97, 4291–6.

37. Mihara, M., Erster, S., Zaika, A., Petrenko, O., Chittenden, T., Pancoska, P. & Moll, U. M. (2003) p53 has a direct apoptogenic role at the mitochondria, Molecular cell. 11, 577–90.

38. Wang, F., Marshall, C. B., Yamamoto, K., Li, G. Y., Plevin, M. J., You, H., Mak, T. W. & Ikura, M. (2008) Biochemical and structural characterization of an intramolecular interaction in FOXO3a and its binding with p53, Journal of molecular biology. 384, 590–603.

39. Wang, F., Marshall, C. B., Yamamoto, K., Li, G. Y., Gasmi-Seabrook, G. M., Okada, H., Mak, T. W. & Ikura, M. (2012) Structures of KIX domain of CBP in complex with two FOXO3a transactivation domains reveal promiscuity and plasticity in coactivator recruitment, Proceedings of the National Academy of Sciences of the United States of America. 109, 6078–83.

40. Lee, C. W., Martinez-Yamout, M. A., Dyson, H. J. & Wright, P. E. (2010) Structure of the p53 transactivation domain in complex with the nuclear receptor coactivator binding domain of CREB binding protein, Biochemistry. 49, 9964–71.

41. Rowell, J. P., Simpson, K. L., Stott, K., Watson, M. & Thomas, J. O. (2012) HMGB1-facilitated p53 DNA binding occurs via HMG-Box/p53 transactivation domain interaction, regulated by the acidic tail, Structure (London, England : 1993). 20, 2014–24.

42. Bochkareva, E., Kaustov, L., Ayed, A., Yi, G. S., Lu, Y., Pineda-Lucena, A., Liao, J. C., Okorokov, A. L., Milner, J., Arrowsmith, C. H. & Bochkarev, A. (2005) Single-stranded DNA mimicry in the p53 transactivation domain interaction with replication protein A, Proceedings of the National Academy of Sciences of the United States of America. 102, 15412–7.

43. Chang, J., Kim, D. H., Lee, S. W., Choi, K. Y. & Sung, Y. C. (1995) Transactivation ability of p53 transcriptional activation domain is directly related to the binding affinity to TATA-binding protein, The Journal of biological chemistry. 270, 25014–9.

44. Chi, S. W., Lee, S. H., Kim, D. H., Ahn, M. J., Kim, J. S., Woo, J. Y., Torizawa, T., Kainosho, M. & Han, K. H. (2005) Structural details on mdm2-p53 interaction, The Journal of biological chemistry. 280, 38795–802.

45. Kussie, P. H., Gorina, S., Marechal, V., Elenbaas, B., Moreau, J., Levine, A. J. & Pavletich, N. P. (1996) Structure of the MDM2 oncoprotein bound to the p53 tumor suppressor transactivation domain, Science (New York, NY). 274, 948–53.

46. Feng, H., Jenkins, L. M., Durell, S. R., Hayashi, R., Mazur, S. J., Cherry, S., Tropea, J. E., Miller, M., Wlodawer, A., Appella, E. & Bai, Y. (2009) Structural basis for p300 Taz2-p53 TAD1 binding and modulation by phosphorylation, Structure (London, England : 1993). 17, 202–10.

47. Follis, A. V., Llambi, F., Ou, L., Baran, K., Green, D. R. & Kriwacki, R. W. (2014) The DNA-binding domain mediates both nuclear and cytosolic functions of p53, Nature structural & molecular biology. 21, 535–43.

48. Weigelt, J., Climent, I., Dahlman-Wright, K. & Wikström, M. (2000) 1H, 13C and 15N resonance assignments of the DNA binding domain of the human forkhead transcription factor AFX, Journal of biomolecular NMR. 17, 181–2.

49. Kim, J., Ahn, D. & Park, C.-J. (2021) FOXO4 Transactivation Domain Interaction with Forkhead DNA Binding Domain and Effect on Selective DNA Recognition for Transcription Initiation, Journal of molecular biology. 433, 166808.

50. Brady, C. A., Jiang, D., Mello, S. S., Johnson, T. M., Jarvis, L. A., Kozak, M. M., Kenzelmann Broz, D., Basak, S., Park, E. J., McLaughlin, M. E., Karnezis, A. N. & Attardi, L. D. (2011) Distinct p53 transcriptional programs dictate acute DNA-damage responses and tumor suppression, Cell. 145, 571–83.

51. Lee, M. S., Lim, K., Lee, M. K. & Chi, S. W. (2018) Structural Basis for the Interaction between p53 Transactivation Domain and the Mediator Subunit MED25, Molecules (Basel, Switzerland). 23.

52. Krois, A. S., Dyson, H. J. & Wright, P. E. (2018) Long-range regulation of p53 DNA binding by its intrinsically disordered N-terminal transactivation domain, Proceedings of the National Academy of Sciences of the United States of America. 115, E11302–e11310.

53. el-Deiry, W. S., Kern, S. E., Pietenpol, J. A., Kinzler, K. W. & Vogelstein, B. (1992) Definition of a consensus binding site for p53, Nature genetics. 1, 45–9.

54. Petros, A. M., Gunasekera, A., Xu, N., Olejniczak, E. T. & Fesik, S. W. (2004) Defining the p53 DNA-binding domain/Bcl-x(L)-binding interface using NMR, FEBS letters. 559, 171–4.

55. Rudiger, S., Freund, S. M., Veprintsev, D. B. & Fersht, A. R. (2002) CRINEPT-TROSY NMR reveals p53 core domain bound in an unfolded form to the chaperone Hsp90, Proceedings of the National Academy of Sciences of the United States of America. 99, 11085–90.

56. Wong, K. B., DeDecker, B. S., Freund, S. M., Proctor, M. R., Bycroft, M. & Fersht, A. R. (1999) Hot-spot mutants of p53 core domain evince characteristic local structural changes, Proceedings of the National Academy of Sciences of the United States of America. 96, 8438–42.

57. Cañadillas, J. M., Tidow, H., Freund, S. M., Rutherford, T. J., Ang, H. C. & Fersht, A. R. (2006) Solution structure of p53 core domain: structural basis for its instability, Proceedings of the National Academy of Sciences of the United States of America. 103, 2109–14.

58. Petty, T. J., Emamzadah, S., Costantino, L., Petkova, I., Stavridi, E. S., Saven, J. G., Vauthey, E. & Halazonetis, T. D. (2011) An induced fit mechanism regulates p53 DNA binding kinetics to confer sequence specificity, The EMBO journal. 30, 2167–76.

59. He, F., Borcherds, W., Song, T., Wei, X., Das, M., Chen, L., Daughdrill, G. W. & Chen, J. (2019) Interaction between p53 N terminus and core domain regulates specific and nonspecific DNA binding, Proceedings of the National Academy of Sciences of the United States of America. 116, 8859–8868.

60. Chien, P. & Gierasch, L. M. (2014) Challenges and dreams: physics of weak interactions essential to life, Molecular biology of the cell. 25, 3474–7.

61. Rabouille, C. & Alberti, S. (2017) Cell adaptation upon stress: the emerging role of membrane-less compartments, Current opinion in cell biology. 47, 34–42.

62. Sukenik, S., Ren, P. & Gruebele, M. (2017) Weak protein-protein interactions in live cells are quantified by cell-volume modulation, Proceedings of the National Academy of Sciences of the United States of America. 114, 6776–6781.

63. Hernandez-Munain, C., Roberts, J. L. & Krangel, M. S. (1998) Cooperation among multiple transcription factors is required for access to minimal T-cell receptor alpha-enhancer chromatin in vivo, Molecular and cellular biology. 18, 3223–33.

64. Nie, Y., Shu, C. & Sun, X. (2020) Cooperative binding of transcription factors in the human genome, Genomics. 112, 3427–3434.

65. Calissi, G., Lam, E. W. & Link, W. (2020) Therapeutic strategies targeting FOXO transcription factors, Nature reviews Drug discovery.

66. Langlois, C., Del Gatto, A., Arseneault, G., Lafrance-Vanasse, J., De Simone, M., Morse, T., de Paola, I., Lussier-Price, M., Legault, P., Pedone, C., Zaccaro, L. & Omichinski, J. G. (2012) Structure-based design of a potent artificial transactivation domain based on p53, Journal of the American Chemical Society. 134, 1715–23.

67. Lee, W., Tonelli, M. & Markley, J. L. (2015) NMRFAM-SPARKY: enhanced software for biomolecular NMR spectroscopy, Bioinformatics (Oxford, England). 31, 1325–7.

